# Unraveling Unbreakable Hairpins: Characterizing RNA secondary structures that are persistent after dinucleotide shuffling

**DOI:** 10.1101/2023.11.30.569461

**Authors:** Alyssa A Pratt, David A Hendrix

## Abstract

The sequence of nucleotides that make up an RNA determines its structure, which determines its function. The RNA hairpin, also known as a stem-loop, is a ubiquitous and fundamental feature of RNA secondary structure. A common method of randomizing an RNA sequence is dinucleotide shuffling with the Altschul-Erickson algorithm, which preserves the dinucleotide content of the sequence. This algorithm generates randomized sequences by sampling Eulerian paths through the de Bruijn graph representation of the original sequence. We identified a subset of RNA hairpins in the bpRNA-1m meta-database that always form hairpins after repeated application of dinucleotide shuffling. We investigated these “unbreakable hairpins” and found several common properties. First, we found that unbreakable hairpins had on average similar folding energies compared to other hairpins of similar lengths, although they frequently contained ultra-stable hairpin loops. We found that they tend to be split by purines and pyrimidines on opposite strands of the stem. Furthermore, we found that this specific sequence feature restricts the number of distinct Eulerian paths through their de Bruijn graph representation, resulting in a small number of distinguishable dinucleotide-shuffled sequences. Beyond this algorithmic means of identification, these distinct sequences may have biological significance because we found that a significant percentage occur in a specific location of 16S ribosomal RNAs. Finally, we present a formula to calculate the number of possible unique dinucleotide shuffled sequences for an input RNA sequence, which has utility for the general application of the Altschul-Erickson algorithm.

## Introduction

RNA has critical roles in translation, gene regulation, genetic sequence encoding, and scaffolding for molecular complexes. Typically, the structure of RNA directs its molecular function, and it is valuable to predict how mutations will disrupt structure. While RNA has a three-dimension tertiary structure, it is often studied with a two-dimensional projection known as the secondary structure. Several approaches have been developed to predict RNA secondary structure including thermodynamic models (Lorenz et al., 2011), machine learning methods including conditional log-linear models (Do et al., 2008), deep neural networks (Chen et al., 2020; Saman Booy et al., 2022), and attention networks (Wang et al., 2020). RNA has several recognizable features that can be predicted by software. In addition to structure prediction, methods have been developed to automatically annotate secondary structural features including hairpin loops, multi-loops, internal loops, bulges, and stems (Danaee et al., 2018).

RNA hairpins, also known as stem-loops, are fundamental features because they are both common and essential, protect messenger RNAs, guide the molecule’s tertiary structure, and serve as recognition sites for proteins (Svoboda & Cara, 2006). RNA hairpins are characterized by two paired strands connected by an unpaired loop. Their diversity in stem length, loop length, and sequence allows for a high specificity in interactions with proteins, especially given their propensity to occur on most types of RNAs in different positions, each of which may serve a different function. For instance, the hairpins of 16S ribosomal RNA tend to have loops of length four (tetraloops), which may contribute to ribosomal stability and protein content depending on their abundance and adjacent base-pairs (Wolters, 1992).

When analyzing RNA, it is common to randomly permute or “shuffle” the RNA sequence, whereby the positions of nucleotides, dinucleotides, or k-mers are rearranged through various algorithms. Shuffled RNA is expected to be free of its original secondary structure, only resembling the original sequence in mononucleotide or dinucleotide frequencies, depending on the method used. Dinucleotide shuffling, often preferred for its ability to retain the original dinucleotide content, has been known as a useful and minimally biased method to create control sequences in bioinformatics for over 40 years (Fitch, 1983). It was found by Clote et al. that the dinucleotide-shuffled versions of functional RNA sequences had a higher predicted folding energy (negative energy being more stable) than the original sequence (with the exception of mRNA), validating the use of a comparison to shuffled sequences to identify functional features (Clote et al., 2005). Dinucleotide shuffling carried out via the Altschul-Erickson algorithm can prevent the control from having a different proportion of thermodynamically favorable pairs (Altschul & Erickson, 1985). The Altschul-Erickson algorithm constructs directed connections between nodes that represent each RNA nucleotide and generates a shuffle by walking along these connections through an Eulerian path from the first nucleotide to the last.

Workman and Krogh found similar results using a heuristic to approximate dinucleotide shuffling, while mononucleotide shuffling was not reliable for the detection of functional RNAs based on different folding energies (Workman & Krogh, 1999). This is dictated by the mechanics of base pairing, for which the energy contributed by a base pair is dependent on the pair it is stacked on. Due to these findings and others, several software tools and analyses have been built around the use of dinucleotide shuffling as a control (Washietl & L. Hofacker, 2007). This allows for a statistical comparison between the folding energy of a sequence and its shuffled variations, such that a statistically significant difference confers the presence of functional RNA. More recently, the ScanFold software tool has used Altschul-Erickson dinucleotide shuffling to detect functional regions in Zika virus, HIV-1, and SARS-CoV-2 RNA (Andrews et al., 2018).

Very little is known about the effectiveness of generating a random sequence (and breaking RNA secondary structures) using dinucleotide shuffling as a control method, even though this is often a base assumption for this work detecting functional RNAs. In the work of Babak et al., it was found that tools used to detect functional RNA elements produced widely different results when applied to a standardized test set (Babak et al., 2007). Though this research utilized a different dinucleotide shuffling algorithm, low complexity sequences that were preserved under shuffling had a large impact on the detection of functional RNAs. This raises questions that have yet to be addressed by current research: are there RNA sequences and structures that persist upon dinucleotide shuffling? Furthermore, how much variation in sequence can be formed from dinucleotide shuffling?

Here we present a subset of hairpins that we call “unbreakable hairpins”, which were found to retain their structure despite repeated Altschul-Erickson shuffling, as well as a formula and computational tool for calculating unique sequence variants that can be created by dinucleotide shuffling. Furthermore, we assess whether there is any biological significance to these unbreakable hairpins.

## Results

We examined the 708,144 hairpins in the bpRNA-1m meta-database; we performed dinucleotide shuffling and predicted secondary structure for each shuffled version. We sought to determine how many shuffles it takes for a sequence to no longer be predicted to form a hairpin structure. Figure 1A shows the number of hairpin-forming sequences that continue to form hairpin structures upon repeated application of the dinucleotide shuffling algorithm (Altschul & Erickson, 1985). We identified 2,364 hairpin-forming sequences that were predicted to form hairpins for each of 1000 shuffled variants, hereafter “unbreakable hairpins”. We sought to characterize the properties of this subset of sequences to explain why they always form hairpins, regardless of how many times the dinucleotide shuffling is algorithm is applied.

**Figure 1:**
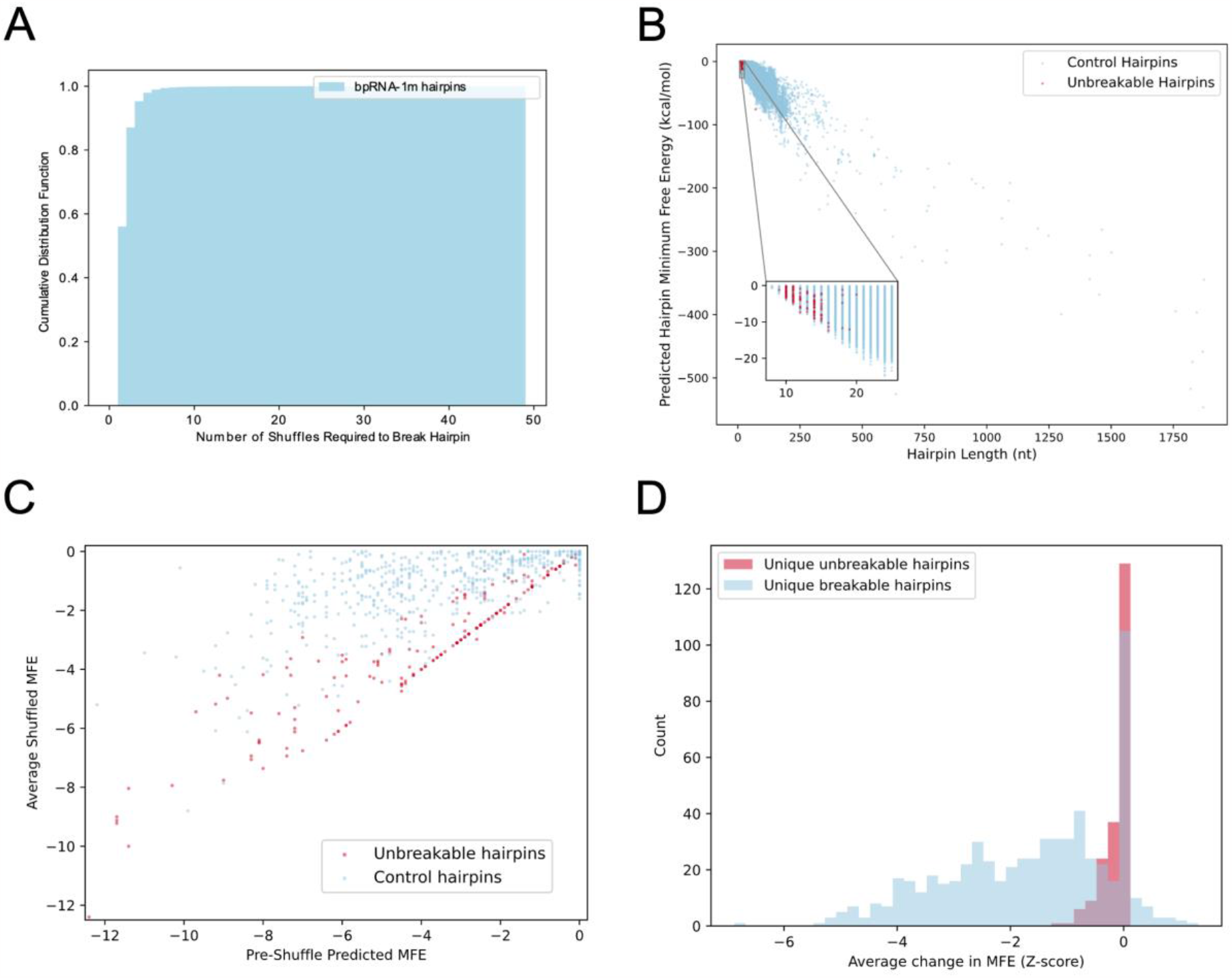
**A**. The cumulative distribution function describing the number of dinucleotide shuffles applied to hairpin sequences such that the shuffled sequence will result in a non-hairpin sequence by secondary structure prediction. **B**. A scatterplot comparing unbreakable hairpins to control hairpins, such that each dot corresponds to a hairpin sequence, the x-axis is the hairpin length, and the y-axis is the predicted minimum free energy of the structure. **C**. A scatterplot showing the average of five minimum free energy values (in kcal/mol) of unbreakable and breakable hairpins before and Altschul-Erickson dinucleotide shuffling, where the diagonal represents no change in hairpin minimum free energy. **D**. A representation of the same results as C, shown as a histogram representing the change in minimum free energy as a Z-score.

### Hairpin Length Distribution

First, we examined the length distribution of the unbreakable hairpins. We observed that the length distribution for the unbreakable hairpin sequences is shorter than the global distribution of hairpin sequences (Figure 1B). We next examined the predicted minimum free energy of the unbreakable hairpins compared to a sample from the global set that has the same length distribution. We observed that the unbreakable hairpins have similar energy values compared to other hairpin sequences of the same length distribution (Figure 1B inset).

Upon shuffling, unbreakable hairpins appeared to have more resistance to shuffling in regard to the estimated minimum free energy structure, while hairpins of a similar length distribution were likely to lose structure, or lose thermodynamic stability, on average across five Altschul-Erickson dinucleotide shuffles (Figure 1C). Many methods use a z-score to quantify the degree of change in free energy upon dinucleotide shuffling (Washietl & L. Hofacker, 2007). We found that unbreakable hairpins have distribution of much higher z-scores, indicating less change upon shuffling (Figure 1D).

We next investigated the length distribution of the hairpin loop itself, defined as the unpaired region, as well as the closing base pair distribution. We found that the unbreakable hairpin loops were predominately of length 4 nt (Figure 2), similar to the global distribution; the favored closing base pair was found to be C:G, which was previously observed to be the most common for tetraloops (Danaee et al., 2018).

**Figure 2:**
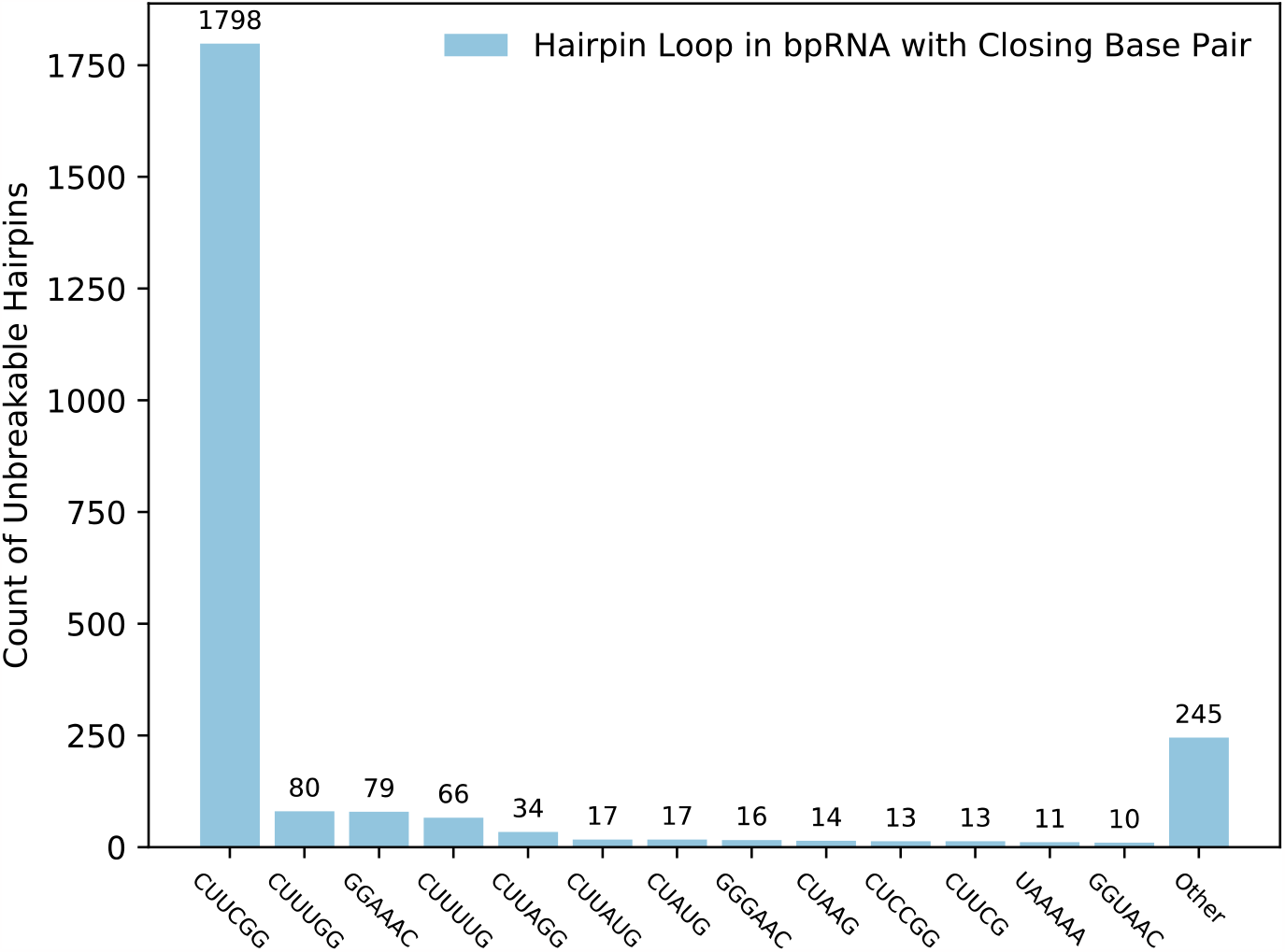
A bar plot representing the number of unbreakable hairpins sharing loop sequences, with the closing base pair included. The most common sequence is C[UUCG]G.

### Hairpin Stem and Loop Sequence Content

We next observed the actual hairpin loop sequence and found it to be enriched with the sequence CUUCGG for 75.2% (1814/2413) of the unbreakable hairpin loops in bpRNA-1m (Figure 2). The CUUCGG tetraloop has been characterized as “ultra-stable” for its experimental stability exceeding thermodynamic predictions to the extent that RNA secondary structure parameters have been modified to account for its additional stability (Tuerk et al., 1988). Additionally, this tetraloop has been found in biologically significant regions of RNA across kingdoms. We found that about 77% of unbreakable hairpins in bpRNA-1m contain this tetraloop, compared to 24% of total hairpins in the database.

We next observed that the unbreakable hairpins tend to be split by purines and pyrimidines. While one strand was typically enriched for either purine (A or G) or pyrimidine (C or U) nucleotides, the other strand had the opposite base composition. 99.9% of unbreakable hairpins were found to exhibit a split on either side of the segment in purines and pyrimidines. The majority of these splits are represented by primarily a cytosine and guanine abundance on each side of the segment, with an average GC abundance of 79% on the segment. GC-rich hairpin segments have been found to exist more frequently than low-GC content hairpins, which is attributed to the strength of the GC-pair (Chan et al., 2009). The specific reflection of abundance as a purine-pyrimidine split, however, is a more specific property. Furthermore, we found that the number of dinucleotide shuffles required to “break” a hairpin—producing a shuffled sequence that results in a non-hairpin structure—was inversely related to the number of “split” dinucleotides that contain both a purine and a pyrimidine (Supplementary Figure 1). Related to this, we observed that on average, the unbreakable hairpins had a lower sequence complexity than breakable hairpins, and unbreakable hairpins that lack the characteristic features of the majority of unbreakable hairpins (Supplementary Figure 2), using Wootton-Federhen sequence complexity (Wootton & Federhen, 1993).

### Hairpin Structure Distribution

Next, we examined whether the unbreakable hairpins are enriched in a particular type of RNA structure found in the bpRNA-1m meta-database, which primarily contains noncoding RNAs. We found that they were enriched for occurrence within 16S ribosomal RNAs (rRNAs) and within the same location. 1,622 out of the 2,364 (68.6%) are found in 16S ribosomal RNA sequences and are identical in sequence.

### Unbreakable Hairpin Identifying Properties

The unbreakable hairpins tend to represent the intersection of three unique properties: the presence of an ultra-stable tetraloop sequence, the presence of a purine & pyrimidine split on either side of the hairpin (Figure 3A). We also observed that the most common location of unbreakable hairpins in the bpRNA-1m meta-database was the same hairpin of 16S ribosomal RNAs, typically hairpin 22 or 23 (Figure 3B).

**Figure 3:**
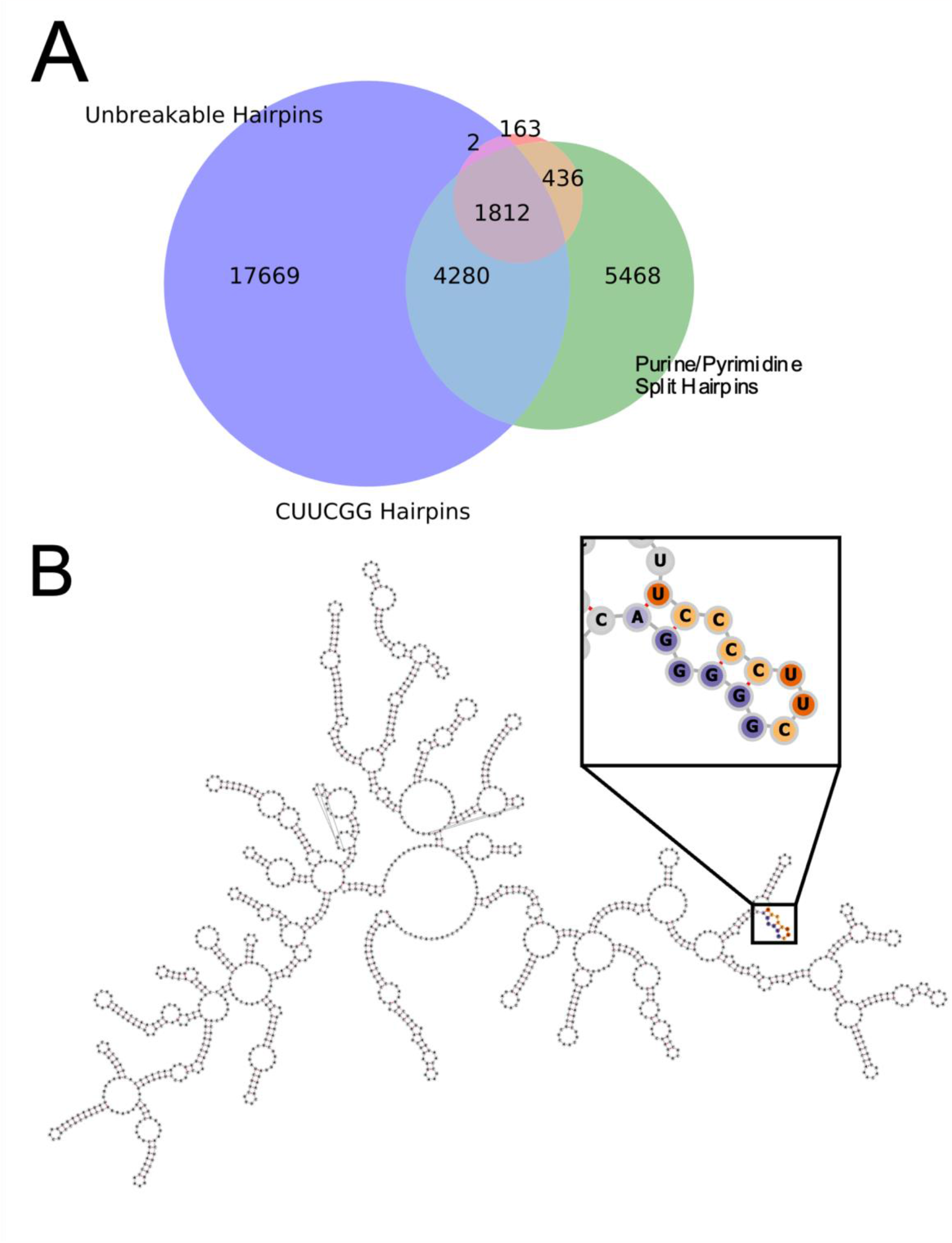
**A:** A venn diagram representation of the shared features of RNA hairpins. The majority of unbreakable RNA hairpins contain a C[UUCG]G loop and split in purines and pyrimidines. **B:** The majority of unbreakable hairpins are located on 16S ribosomal RNA.

### Enumeration of Distinct Sequences Generated by Dinucleotide Shuffling

Finally, we examined the sequences of the unbreakable hairpins in the context of the Altschul-Erickson algorithm. This algorithm generated different dinucleotide-shuffled sequences by selecting random Eulerian circuits within the de Brujin graph for the sequence (Figure 4). The de Bruijn graph *G* = (*E, V*)is such that each nucleotide is represented by a vertex *v*∈*V*, and the edges *E* between them correspond to dinucleotides that occur in the input sequence (e.g. (*A, G*)∈ *E*). To ensure that every vertex has the same number of ingoing and outgoing edges, an additional “false edge” is added between the vertices corresponding to the first and last character. Eulerian circuits are defined as paths through the graph that visit every edge one.

**Figure 4:**
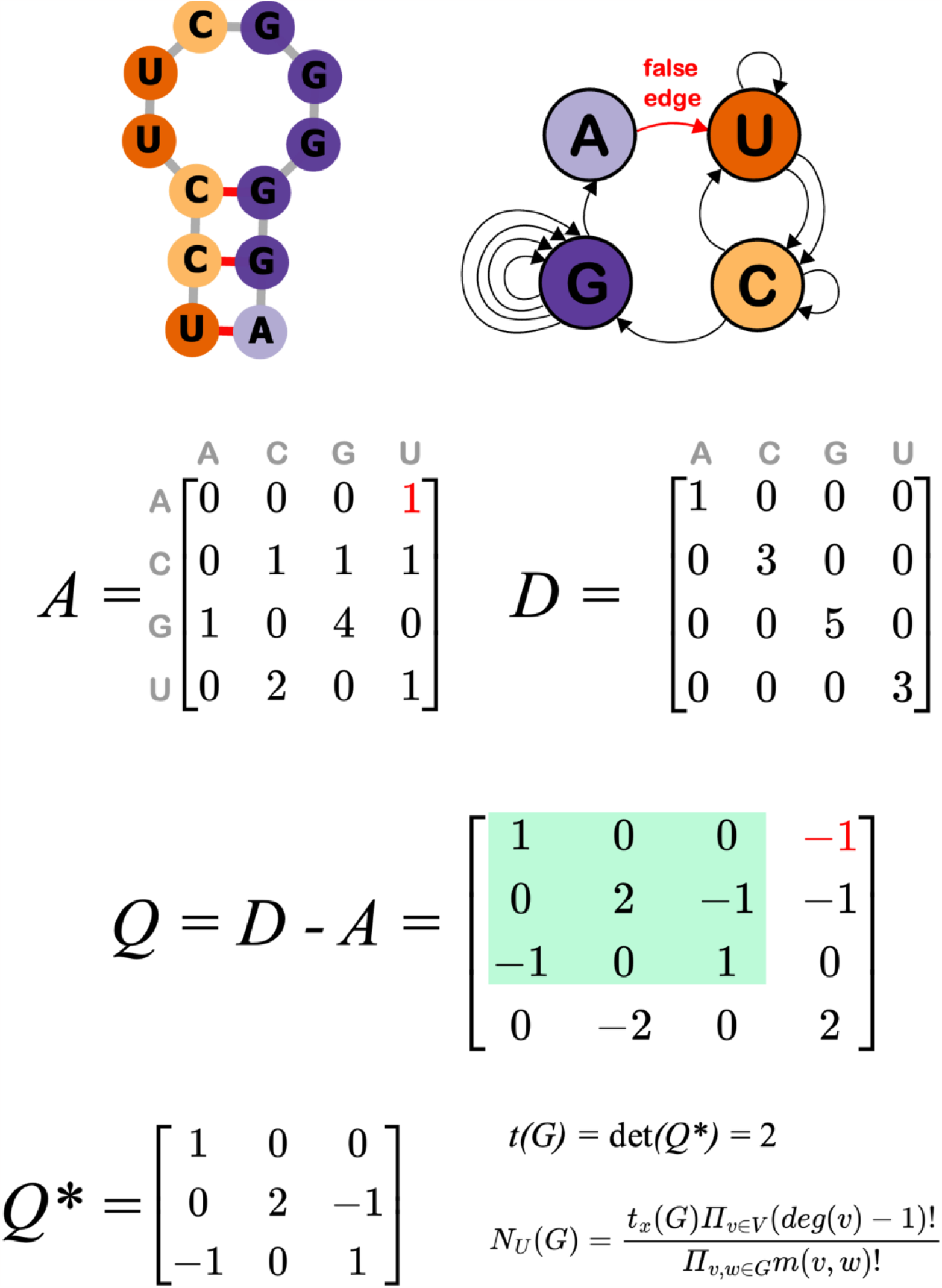
The calculation of the number of unique sequences possible through dinucleotide shuffling. The matrix *A* corresponds to an adjacency matrix that includes the false. The matrix *D* is a diagonal matrix containing the in-degrees of each nucleotide vertex. The matrix Q is the difference of A and D, and Q* is reduced by removing the row and column where the path begins (the nucleotide that the false edge points to. The calculation of the determinant of the resultant *Q\** produces a value that that can be used to calculate *N*_*U*_(*G*).

The number of Eulerian circuits of such a graph can be computed using the de Bruijn, van Aardenne-Ehrenfest, Smith and Tutte (BEST) theorem (van Aardenne-Ehrenfest & de Bruijn, 1951). It states that the number of Eulerian circuits *N*_*E*_ (*G*)is given by the following equation:

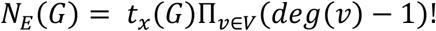

In this expression the term *t*_*x*_(*G*)is the number of arborescences, which are trees directed toward the root from a fixed vertex *x*. This term can be computed as the determinant of the matrix *Q*, defined as the difference of the adjacency matrix *A* and the diagonal *D*. The elements of *Q* are therefore such that *Q*_*ij*_ equals – *m* for a graph with m edges from *i* to *j*, and *Q*_*ii*_ equals the number of ingoing edges minus the number of self-edges (Figure 4D-F). Based on Kirchhoff’s theorem for directed multigraphs, the number of arborescences at a vertex *i* is equal to the determinant of the matrix resulting from the removal of the *i*-th row and column of *Q* (Kirchhoff, 1847). In this case, this results in a 3x3 matrix (Figure 4F).

A dinucleotide shuffled sequence is generated by the series of nucleotides corresponding to the vertices visited in an Eulerian circuit traversing the de Bruijn graph. We illustrate an example of the secondary structure and de Bruijn graph for an unbreakable (Figure 5A,B) and breakable example (Figure 5C,D). While the BEST theorem counts the number of Eulerian circuits, it does not count the number of distinct possible dinucleotide-shuffled sequences because many such sequences are the same, despite arising from different Eulerian circuits. The equation counts degree of the vertices, but while the edges in the graph are distinct, the resulting character sequence may be the same; hence it overcounts the number of unique generated sequences. We therefore consider a modified form of this equation that divides by the total number of rearrangements of the indistinguishable edges, connecting the same nodes. This divisor is given by the product of the factorial of *m*(*v, w*)number of parallel edges in the graph connecting vertices *v* and *w*.

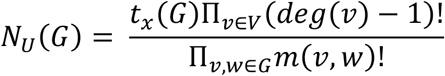

**Figure 5:**
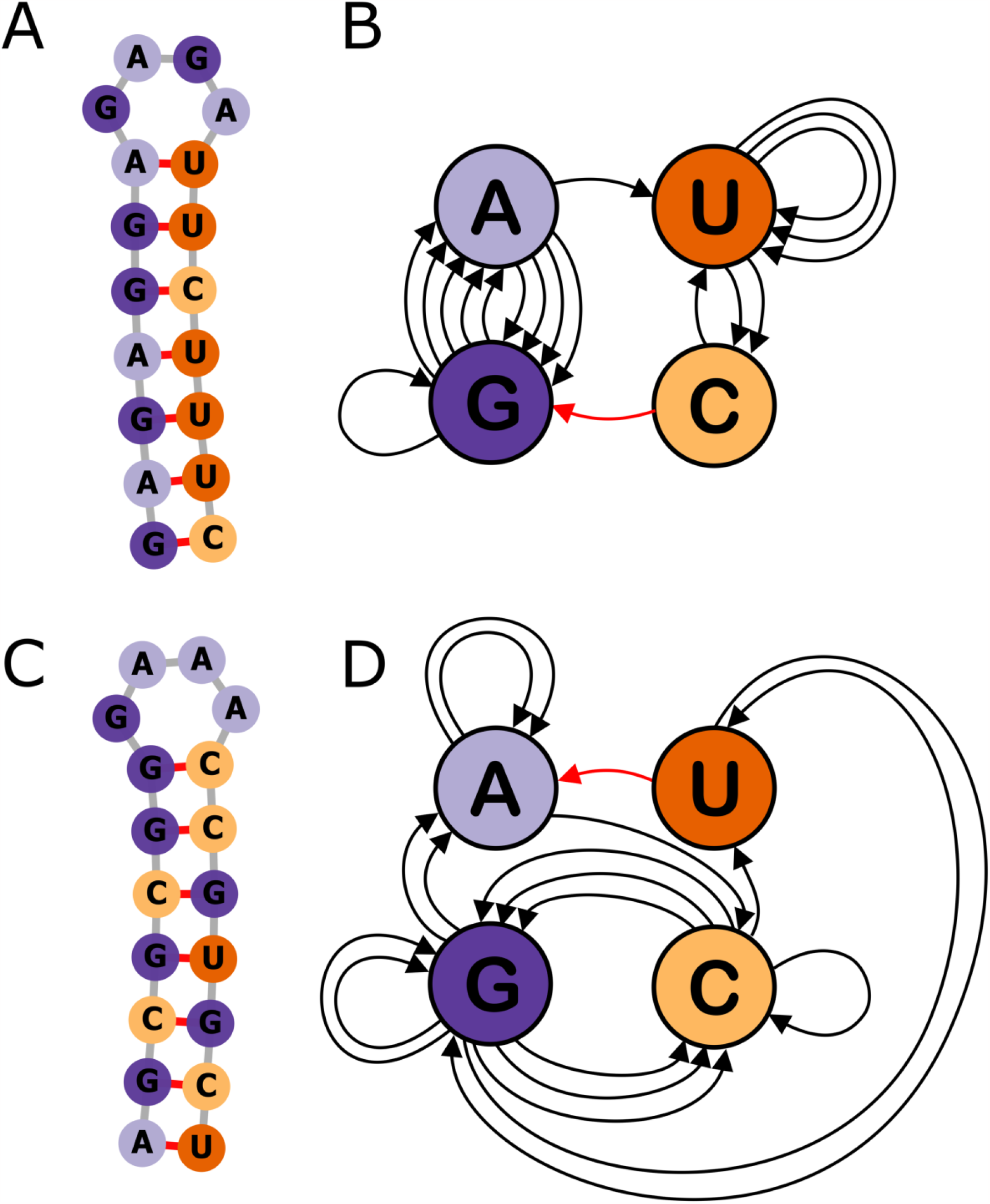
**A**. An example of a secondary structure of an unbreakable hairpin. **B**. The de Bruijn graph representation of the unbreakable hairpin in panel A. **C**. An example of a secondary structure of a breakable hairpin. **D**. The de Bruijn graph representation of the unbreakable hairpin in panel C.

This equation updates equation from the BEST theorem to compute *N*_*U*_ (*G*), the number of unique Eulerian circuits, avoiding what would otherwise be multiple counting of circuits that result in the same output sequence.

### Biological significance of unbreakable hairpins

To elucidate potential biological relevance, we conducted a phylogenetic analysis of 16S sequences that contain unbreakable hairpin sequences in Bacteria. We found that sequences in the class Bacilli were more likely to contain unbreakable hairpins than those in other classes Supplementary Figure 3. Within Bacilli, the distribution among orders, families, and genera was unequal. For example, at the order-level, less than 1% of sequences in Lactobacillales contained unbreakable hairpin sequences as opposed to approximately 25% of sequences in Bacillales. Within Bacillales, 85% of Sporolactobacillacea sequences contained unbreakable hairpins, indicating a potential connection to endospore formation.

To determine whether there is a connection between endospore formation and unbreakable hairpins, we identified the proportion of sequences in each genus within Bacilli and Alphaproteobacteria, which both contained some unbreakable sequences, and compared this to whether or not the genera were known to form endospores. Approximately 27.3% of genera were not found to have sufficient research to support endospore formation or a lack thereof. With the remaining genera, a one-tailed t-test for independence produced a p-value of < .01. A Fisher contingency table p-value was also <.01, supporting the positive association between endospore formation and unbreakable sequences.

Finally, we found that the sequence features that characterize unbreakable hairpins may also confer a robustness to sequence mutations. We observed that unbreakable hairpins on average show less of a change in stem length when a random single-nucleotide insertion or deletion is introduced (Supplementary Figures 4 and 5). This trend is strongest for deletions, but in both cases a greater proportion of unbreakable hairpins are unchanged in length.

## Discussion

Dinucleotide shuffling is widely used to generate control sequences in the analysis of RNA. However, this method does not account for the varying potential of dinucleotide shuffling to produce distinct outputs across RNA sequences, which may produce results that are biased towards structures that are “broken” upon shuffling. Counting the number of possible unique sequences that can be generated through dinucleotide shuffling has significant implications for bioinformatics work with RNA sequences. Here we presented a method to quantify the number of potential distinct dinucleotide shuffled sequences, given by the equation *N*_*U*_(*G*). Calculating *N*_*U*_(*G*)may be used in the software described above to provide a score quantifying the potential for structures to have little difference between original and shuffled free energies. As illustrated in Figure 6, it is unlikely that more complex RNA features may have the unbreakable property due to the necessity of 1 or fewer transitions between purine and pyrimidine nucleotides. An exception shown is the H-type pseudoknot, which could contain a single transition, though there could be no other structure between the closing pair of the hairpin stem and the nucleotides paired to the loop portion of the hairpin, so structure possibilities are limited. An example of a simple H-type pseudoknot with one transition is shown in Figure 6B.

**Figure 6:**
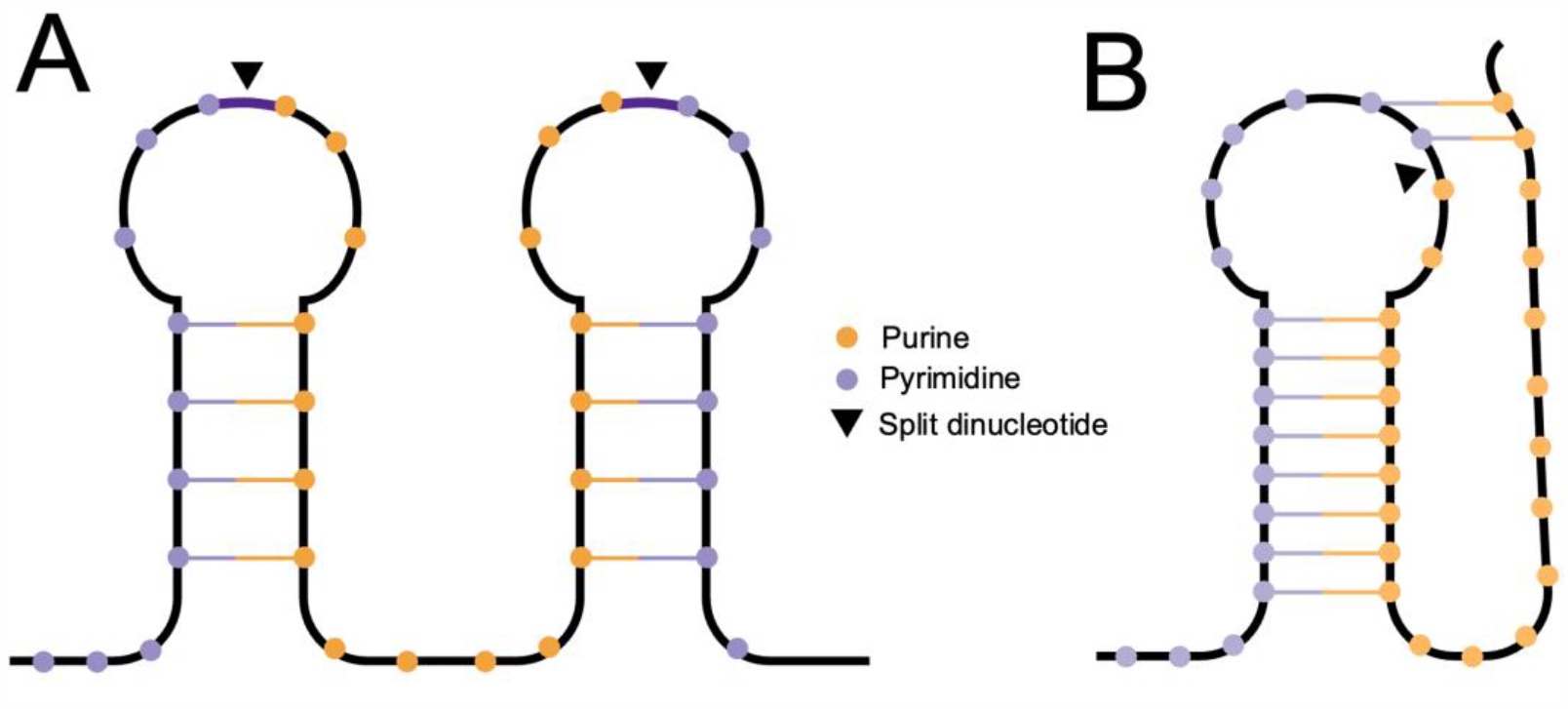
***A***. The structure of unbreakable hairpins prevents multiple hairpins from forming an unbreakable structure, as multiple purine/pyrimidine splits must be present. **B**. It is feasible for a single H-type RNA pseudoknot to be unbreakable if there are no further splits.

It remains unclear whether unbreakable hairpins are an oddity of the dinucleotide shuffling algorithm, or if they are biologically relevant. As the majority of unbreakable hairpins were located on in a specific region within 16S ribosomal RNAs, where the structures of tetraloop hairpins are known to facilitate interactions that enhance protein stability, the unbreakable features of the hairpin may allow for sustained stability despite deletions (Wolters, 1992). Since unbreakable hairpins have a lower sequence complexity than breakable hairpins of similar length, and the majority have only one “split” dinucleotide with both a purine and a pyrimidine, single-nucleotide or dinucleotide deletions are less likely to affect the secondary structure than in a control hairpin of similar length. We have shown a statistical relationship between unbreakable hairpins and endospore-forming bacteria in the class Bacilli. Unbreakable hairpin sequences may confer a robustness due to their retained structure upon rearrangement, which could be valuable to bacterial endospores which are known to retain stability upon exposure to radiation, desiccation, and heat (Setlow, 2006).

In RNA tertiary structures, it has been found that a split in purines and pyrimidines is associated with Watson-Crick base pairing in the MALAT1 long noncoding RNA nuclear retention element. Brown et. al. described a preference for pyrimidine-purine pairs on this triple helix over purine-pyrimidine pairs, as determined by chemical probing using Watson-Crick base pair replacement mutations (Brown et al., 2014). All base pairs in the triple-helical region retaining a purine-pyrimidine split in the wild-type form. This supports the hypothesis of purine-pyrimidine splits increasing the robustness of RNAs.

UUCG loops are currently considered an “experimental tool” that can stabilize the secondary structure of RNAs without affecting interactions, as it has not been found to bind to proteins or contribute to RNA:RNA interactions (Hall, 2015). Since the initial categorization 16S rRNA tetraloops, research has found that the ultra-stable UNCG hairpin loops, and particularly those containing the sequence C(UUCG)G, with the C and G outside the parentheses indicating the closing base pair, are more abundant than their thermodynamic stability and base pair frequency would indicate (Hall, 2015; Tuerk et al., 1988; Varani, 1995). Suggested roles of UNCG hairpins in rRNA and other locations include interactions with ribosomal proteins, forming nucleation sites for RNA folding, and protection against degradation by nucleases (Thapar et al., 2014; Varani, 1995). Indeed, UNCG hairpins have been found to serve as sites of zinc finger protein interactions, such as the Rous Sarcoma Virus packaging signal’s UGCG tetraloop, which interacts with a zinc knuckle on the nucleocapsid protein and appears to affect viral assembly without any conformational change in the RNA (Thapar et al., 2014).

Despite the ubiquity of C(UUCG)G loops, no such specific functional interaction has been found these tetraloops. Experimental studies have confirmed that proteins such as the double-stranded RNA-binding protein, which bind any dsRNA of sufficient length, will bind a C(UUCG)G tetraloop, but do not appear to have specific intermolecular interactions (Ramos et al., 2000). Despite its highly stable form, the structure is also nonrigid as found by nuclear magnetic resonance imaging (Bottaro et al., 2020). Recent findings suggest that there may be two primary types of canonical tetranucleotide turns that correspond to GNRA- and UNCG-type conformations, called U-turns and Z-turns, which reflect canonical structures that may not always be distinguishable from sequence (D’Ascenzo et al., 2017). Of the three most common unbreakable hairpin loop sequences, UUCG, UUUG, and GAAA, only UUCG corresponds to the typical UNCG tetraloop sequence, but all three sequences match known sequences of Z-turn tetraloop conformations.

## Methods

### Unbreakable Hairpin Identification

Unbreakable hairpins were identified as a subset of all hairpin segments listed in bpRNA-1m database (Danaee et al., 2018), which includes full stems and hairpin loops stored in the FASTA file format. Each segment listed was then processed by RNAfold version 2.1.9 (Lorenz et al., 2011) to determine the dot-bracket sequence of the predicted hairpin structure. To filter against invalid hairpins, hairpin segments that were not comprised of solely A, C, G, and U characters were eliminated, as were segments that did not meet two thresholds to qualify as hairpins: balance and base pair density. Balance scores were computed from the distribution of left and right parenthesis in the dot-bracket sequence, while base pair density scores were computed from the total fraction of left parenthesis. If the balance score exceeded 1.86 and the base pair density exceeded 0.25, hairpins proceeded to a series of Altschul-Erickson dinucleotide shuffles (Deng et al., 2019)

According to the Altschul-Erickson algorithm, each hairpin was shuffled repeatedly until its balance score, or base pair density was no longer within the thresholds. If hairpins could be shuffled at least 1000 times and remain a hairpin, they were stored as unbreakable hairpins.

### Control Hairpin Set

Control hairpins were randomly sampled from the hairpin segments in the bpRNA-1m database that were of similar or equal length to each unbreakable hairpin, creating a comparable set of equal size. If an exact match in length could not be found for each unbreakable hairpin, the control hairpin with the next-closest length was kept. The control and unbreakable hairpin set are of an equal size.

### Hairpin Content

We found the length distribution of unbreakable hairpin sequences as well as the global length distribution of RNA hairpins in bpRNA-1m from their respective FASTA sequences. We then computed the predicted minimum free energy (MFE) of each unbreakable hairpin and control hairpin from RNAfold 2.1.19 using default parameters.

We also investigated the loop of each hairpin using similar methods, as the loop of each hairpin segment is stored individually. We searched for the known ultra-stable sequence motif “CUUCGG” (Tuerk et al., 1988) in each hairpin in the control and unbreakable regions to find evidence of enrichment. We also determined the propensity of a total purine/pyrimidine split in hairpin segments. We defined a “total split” as a hairpin with nucleotides that were entirely purines or pyrimidines on one side of the hairpin loop (one stem) and the other on the other side, disregarding the content of the loop. We recorded the content of each split unbreakable and control hairpin.

### Hairpin Structure Distribution

Using the RNA structure annotation tool, bpRNA, we identified the location of each unbreakable hairpin. Of 2,413 unbreakable hairpins, 1,116 (46%) were the 22^nd^ hairpin in the molecule, and 383 were the 23^rd^ hairpin in the molecule – typically of 16S ribosomal RNA. Overall, 2,064 unbreakable hairpins (86%) were classified as 16S ribosomal RNA in the bpRNA-1m meta-database.

### Mathematical Basis of Dinucleotide Shuffling

We used the de Bruijn, van Aardenne-Ehrenfest, Smith and Tutte (BEST) theorem to calculate the number of unique Eulerian circuits possible for a given sequence, as each individual Eulerian circuit represents a dinucleotide shuffle (van Aardenne-Ehrenfest & de Bruijn, 1951). In order to determine the count of unique *sequences* rather than circuits, we divided our result for each sequence by the product of the factorials of the number of each of its parallel edges, eliminating circuits that differ only by a parallel edge.

## Data Availability

RNA sequences used for this analysis were downloaded from bpRNA-1m, and filtered using CD-HIT with a sequence identity value of 1.0 (Fu et al., 2012). The unbreakable set of RNA hairpins, as well as the control RNA hairpin set used in the analyses, are available at http://github.com/alyssapratt/unbreakable-hairpins. All Python scripts used to generate the figures in this analysis are available within the same GitHub repository.

