## Supplementary Figures for "Unraveling Unbreakable Hairpins: Characterizing RNA secondary structures that are persistent after dinucleotide shuffling"

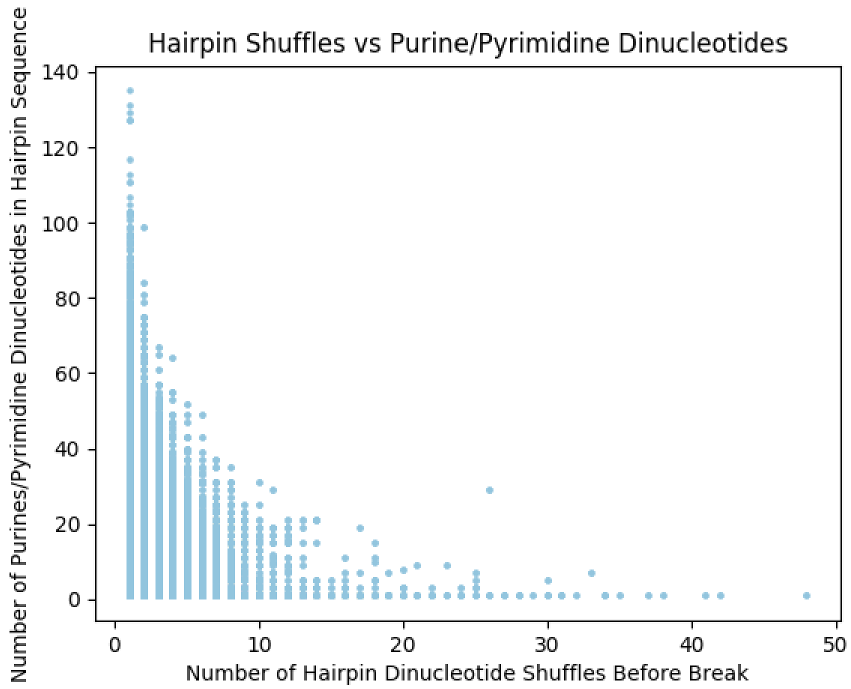


*Supplementary Figure 1*: The number of dincleotide shuffles for each RNA hairpin until a "break", resulting in a non-hairpin structure, has an inverse relationship with the number of purine/pyrimidine split dinucleotides.

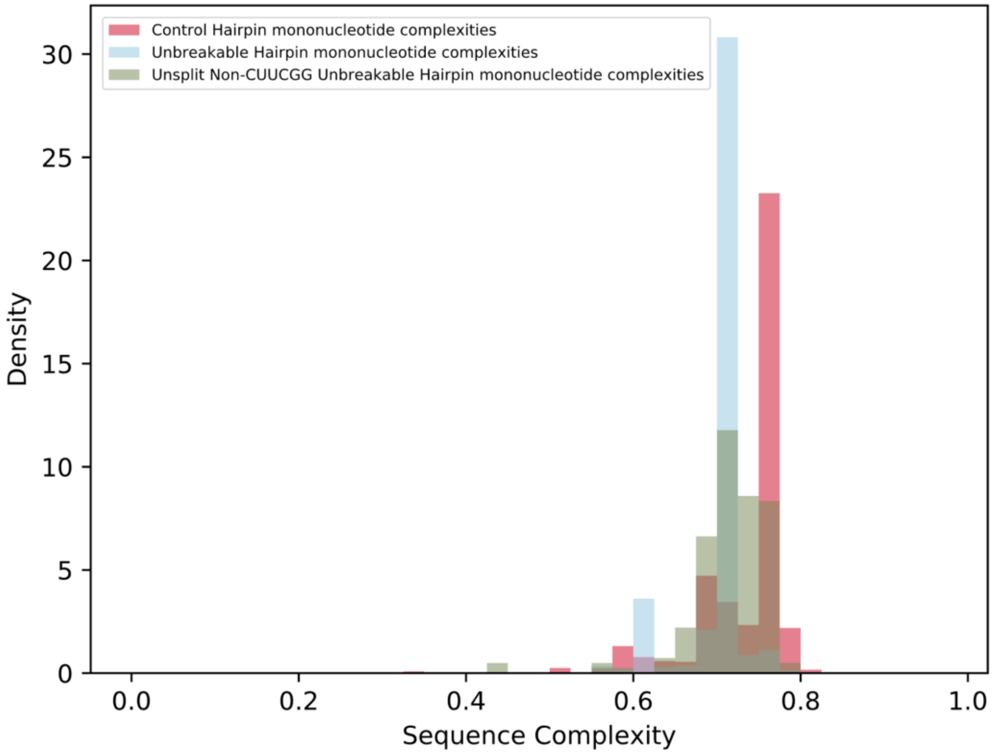


*Supplementary Figure 2*: The sequence complexity of hairpins varies between a representative set of control hairpins, unbreakable hairpins, and hairpins that lack two of the key features of the majority of unbreakable hairpins: a purine/pyrimidine split and C[UUCG]G tetraloop. On average, unbreakable hairpins have lower sequence complexity.

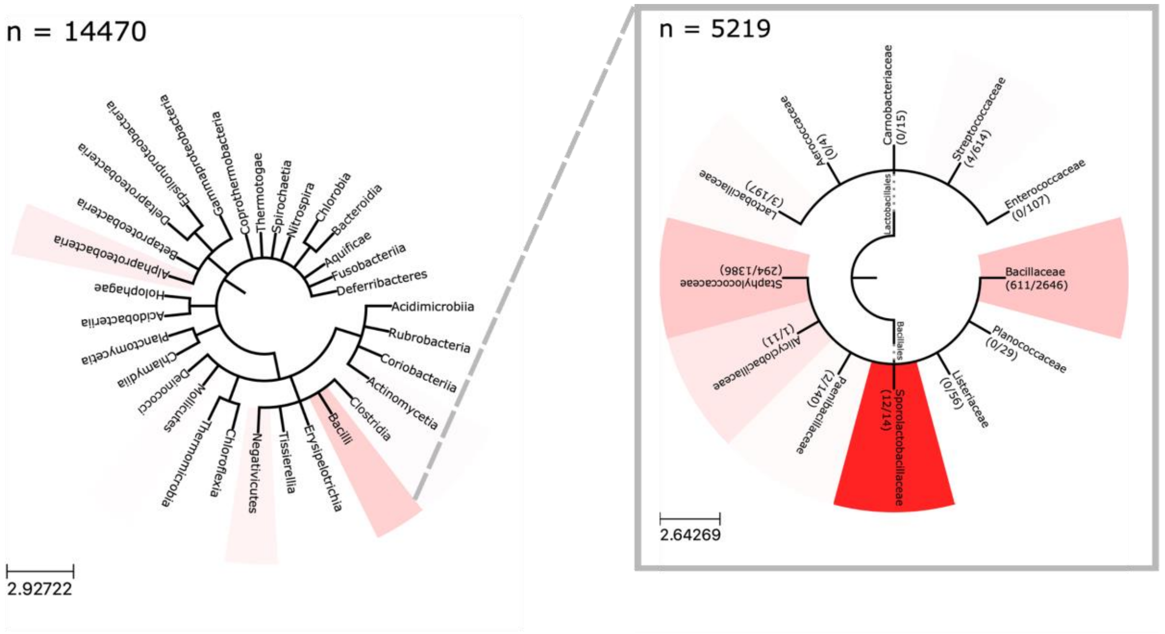


*Supplementary Figure 3*: Phylogenetic distribution of unbreakable hairpins in Bacillales, with white representing no unbreakable hairpins in all sequenced 16S RNA and a complete saturation of red representing 100% of sequences containing unbreakable hairpins. Unbreakable hairpins appear to be enriched in 16S rRNA of Bacillales, particularly in endospore-forming orders.

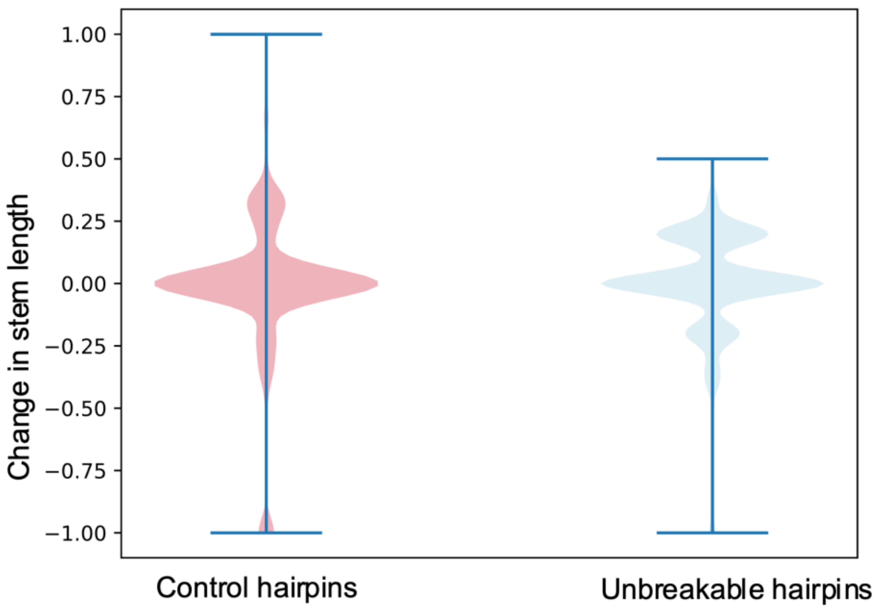


Supplementary Figure 4: Violin plots representing the relative change in stem length for unbreakable hairpins and control hairpins with the same length distribution after a single-nucleotide random insertion. A change of -1 represents a complete loss of hairpin structure.

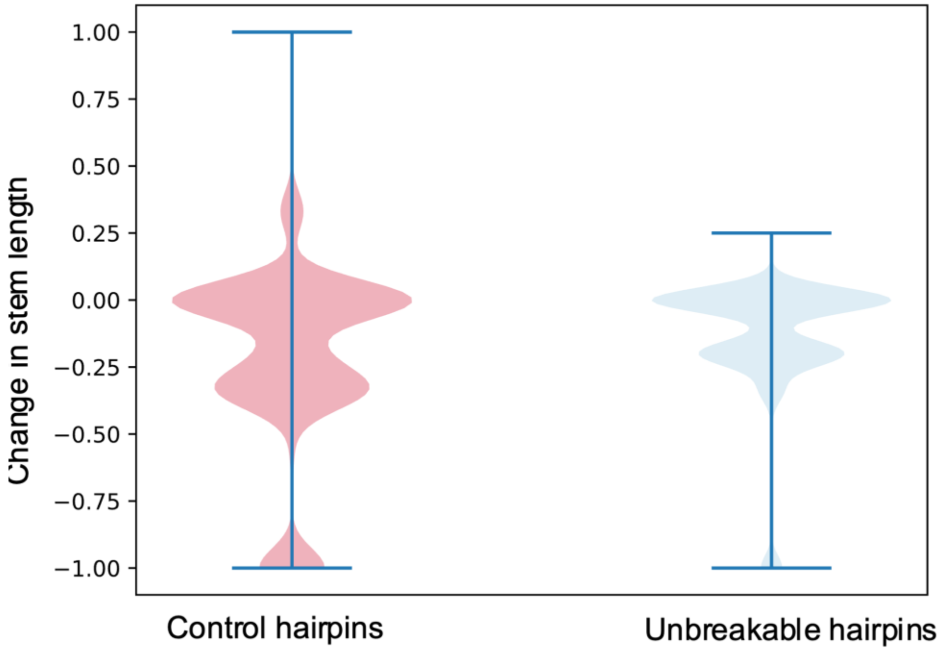


Supplementary Figure 5: Violin plots representing the relative change in stem length for unbreakable hairpins and control hairpins with the same length distribution after a single-nucleotide random deletion. A change of -1 represents a complete loss of hairpin structure.
